# A cell-free assay for rapid screening of inhibitors of hACE2-receptor - SARS-CoV-2-Spike binding

**DOI:** 10.1101/2021.10.06.462907

**Authors:** Nanami Kikuchi, Or Willinger, Naor Granik, Noa Navon, Shanny Ackerman, Ella Samuel, Tomer Antman, Noa Katz, Sarah Goldberg, Roee Amit

## Abstract

We present a cell-free assay for rapid screening of candidate inhibitors of protein binding, focusing on inhibition of the interaction between the SARS-CoV-2 Spike receptor binding domain (RBD) and human angiotensin-converting enzyme 2 (hACE2). The assay has two components: fluorescent polystyrene particles covalently coated with RBD, termed virion-particles (v-particles), and fluorescently-labeled hACE2 (hACE2F) that binds the v-particles. When incubated with an inhibitor, v-particle - hACE2F binding is diminished, resulting in a reduction in the fluorescent signal of bound hACE2F relative to the non-inhibitor control, which can be measured via flow cytometry or fluorescence microscopy. We determine the amount of RBD needed for v-particle preparation, v-particle incubation time with hACE2F, hACE2F detection limit, and specificity of v-particle binding to hACE2F. We measure the dose response of the v-particles to a known inhibitor. Finally, we demonstrate that RNA-hACE2F granules trap v-particles effectively, providing a basis for potential RNA-hACE2F therapeutics.

## INTRODUCTION

The current COVID-19 pandemic, caused by the SARS-CoV-2 virus ^1,2^, has resulted in an unprecedented need for tools that combat the spread of the virus, and for therapeutics for those infected. SARS-CoV-2 virions enters the host cells via interaction between the receptor binding domain of the viral Spike protein (RBD), and hACE2 on the host cell surface ^3,4^. Characterization of both RBD-hACE2 binding and the inhibition of this binding are crucial to three different aspects of controlling the pandemic. First, it is necessary to rapidly quantify the hACE2 binding affinity of naturally-arising RBD mutants in order to assess the impact of these mutants on viral transmission and case load ^5^. Second, inhibition of RBD-hACE2 binding is a useful way to quantify the neutralization capabilities of the immune response and thus provides a metric for assessing SARS-CoV-2 vaccines based on RBD-hACE2 inhibition ^6,7^, whether in inactivated (Sinovac) ^8^, DNA (AstraZeneca) ^9^, mRNA (Pfizer/BioNtech and Moderna) ^10,11^, or protein (Novavax) ^12^ form. Finally, inhibition of RBD-hACE2 interaction could protect healthy host cells at early stages of infection, and is therefore a desired property of candidate therapeutics ^13,14^. For all three aspects, namely characterizing hACE2 mutants, assessing immune response, and quantifying inhibition of candidate therapeutics, an assay that quantifies RBD-hACE2 interaction, and enables rapid identification of small molecules that inhibit RBD-hACE2 interaction, is of great interest.

Repurposing of drugs approved by either the FDA or the EMA is perhaps the fastest path for identification of approved therapeutics for emerging diseases ^15,16^. In silico strategies are currently being employed to identify approved drugs that could be repurposed for COVID-19 ^17^. The standard experimental screen for candidate compounds is an in vitro viability assay ^18^, in which ex vivo cells are first mixed with the compounds, and then infected with the virus. The percentage of viable cells is compared to their percentage in infected+non-treated and non-infected controls. However, high-throughput screening with cell culture requires multiple days, is relatively expensive, and requires Biosafety Level 3 biocontainment conditions. Also, assay results may differ between labs due to differences in cell strain, growth conditions, and inherent variability in biological response. Pseudovirus assays for SARS-CoV-2 inhibitors ^19,20^ require only Biosafety Level 2, but may still suffer from relatively high expense and inherent variability due to the cellular component. These constraints provide motivation for cell-free screening alternatives ^21^.

Ideally, a cell-free assay for screening of inhibitors of protein-protein interaction should satisfy the following requirements: detection using standard lab equipment, repeatability, ease of use, flexibility, and low cost. Since protein sizes are well below the optical diffraction limit, some form of bulk measurement is required. To our knowledge, the only commercial cell-free option currently available for screening RBD-hACE2 inhibitors (Cayman Chemical, Cat. 502050) consists of antibody-coated surface that binds antigen-RBD. Horseradish peroxidase (HRP)-hACE2 is introduced in the presence or absence of an inhibitor candidate. Excess HRP-hACE2 is rinsed, and HRP activity is measured optically at 450 nm via plate reader. However, this assay requires expensive reagents, and multiple washing steps that could affect assay repeatability. In this work, we developed a particle-based fluorescence assay for rapid screening of candidate inhibitors of RBD-hACE2 interaction without the need for live cells or viruses (see Fig. 1A). Our assay utilizes a fluorescent version of hACE2 (containing mCherry, for sequence see Supplementary Table 1), which eliminates the use of antibodies and any related labeling and rinsing steps. We use fluorescent particles covalently-coated with RBD, which we term v-particles, as the surface on which binding occurs. Possible roles of small particles in the context of COVID-19 have been discussed elsewhere ^22–24^. In our assay, the v-particles provide a number of benefits: first, during v-particle preparation, unattached RBD can be removed from the v-particle stock via centrifugation, so that all RBD-binding events occur at the v-particle surface. This could enable production of a ready-to-use product that can be more easily shipped and stored than a coated microplate. Such a ready-to-use product could enable better quality control of the assay and yield more reproducible results ^25^. Second, the v-particles provide a versatile platform: v-particles can be prepared with any choice of viral proteins (e.g., various RBD mutants) or be adapted to display any desired component without necessitating a particular chemical modification (Fig. 1B). Finally, v-particles are large enough to be easily detectable using either flow cytometry or standard fluorescence microscopy, and enable clear distinction of bound hACE2 from unbound hACE2 when assayed via flow cytometer or microscope without the need for cleanup via centrifugation or buffer exchange.

**Figure 1.**
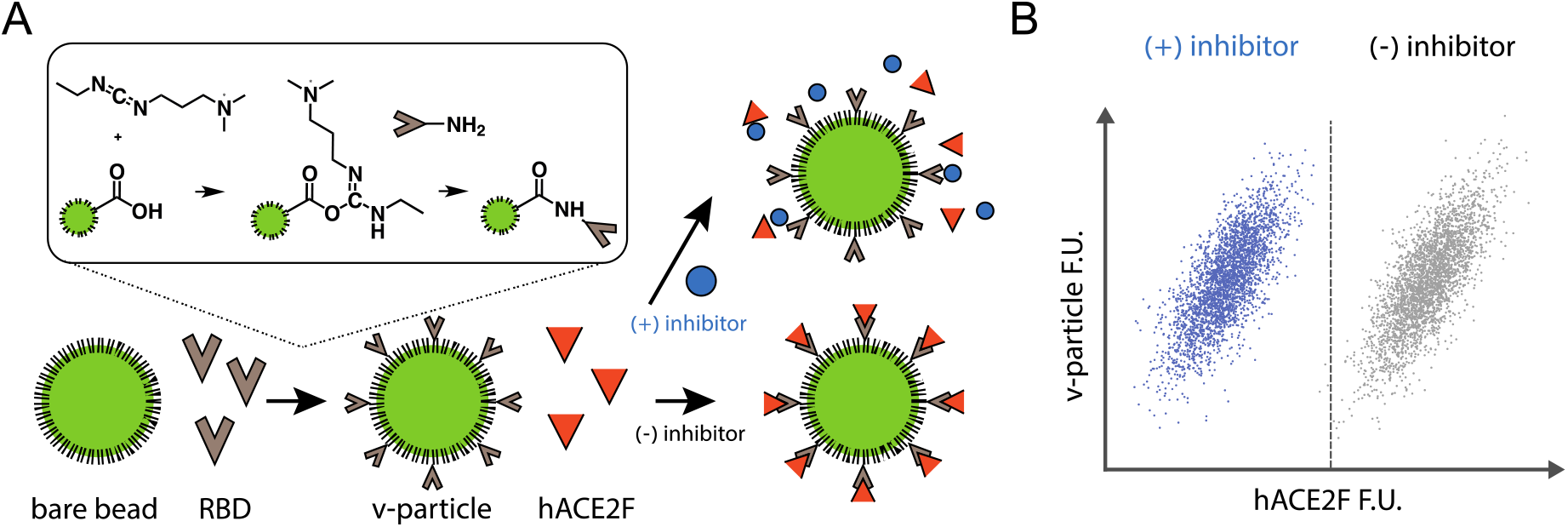
Schematic of the v-particle binding assay. (A) RBD is covalently attached to fluorescent polystyrene particles (green), yielding virion-like particles (v-particles). V-particles are incubated with (red) hACE2F in the presence and absence of a candidate inhibitor. (B) Inhibitor activity is measured by the decrease in the red fluorescence of the v-particles, due to reduced hACE2F binding.

## RESULTS AND DISCUSSION

The RBD for the v-particles was expressed from a plasmid encoding his-tagged RBD that was a gift from the Krammer lab (see sequence in Supplementary Table S1). RBD ^26^ was extracted from Freestyle 293F cells (Thermo Fisher) following the manufacturer’s protocol (for full details, see Supplementary Methods). We note that the RBD contains post-translational modifications ^27^ that are not enzymatically supported in bacterial cells. For v-particle generation, SPHERO carboxyl fluorescent yellow particles with 0.7-0.9 µm diameter were purchased (Spherotech Inc., specified batch diameter was 0.92 µm). The RBD protein was attached to the particles by two-step carbodiimide crosslinker chemistry (see Fig. 1B) using N-(3-Dimethylaminopropyl)-N′-ethylcarbodiimide hydrochloride (EDC, Sigma Aldrich) and N-hydroxysulfosuccinimide sodium salt (Sulfo-NHS, Sigma Aldrich). (See Supplementary Methods for full details.)

For the v-particle binding partner, we expressed and secreted a his-tagged, fluorescently-labeled version of the extracellular domain of hACE2 from HEK 293F cells, similarly to RBD expression (see Supplementary Methods). We note that we were not able to express the fusion hACE2-mCherry in significant titer, and instead resorted to expression of a longer fusion protein, hACE2-mCherry-tdPP7 (see sequence in Supplementary Table S1), which proved to be highly expressed. We refer to hACE2-mCherry-tdPP7 as hACE2F in the following.

We verified the specificity of v-particle binding to hACE2F by comparing to v-particle binding to mCherry (Fig. 2A, for comparison to tdPP7-mCherry see Fig. 2B). We measured v-particle binding to either hACE2F or mCherry, as follows. The results for the specificity assay are shown in Fig. 2A. The plot shows that v-particles incubated with mCherry exhibit a modest and continuous concentration-dependent shift in fluorescence from the non-fluorescent control, that is consistent with non-specific binding. For v-particles incubated with comparable amounts of hACE2F, a fluorescent population of v-particles emerges corresponding to a 2-3 order of magnitude shift in fluorescence for the non-fluorescent controls, indicating specific binding of hACE2F to the RBD displayed on the v-particles. Interestingly, the dose response observed here is a digital-like increase in the fluorescent bead-fraction rather than an analog shift of the v-particle population from no-fluorescence to full-fluorescence values. The v-particles exhibited the same qualitatively-different behavior for hACE2F vs a tdPP7-mCherry control, albeit with smaller population in the high-fluorescence peak for the hACE2F, approximately three months after v-particle preparation (see Fig. 2B).

**Figure 2.**
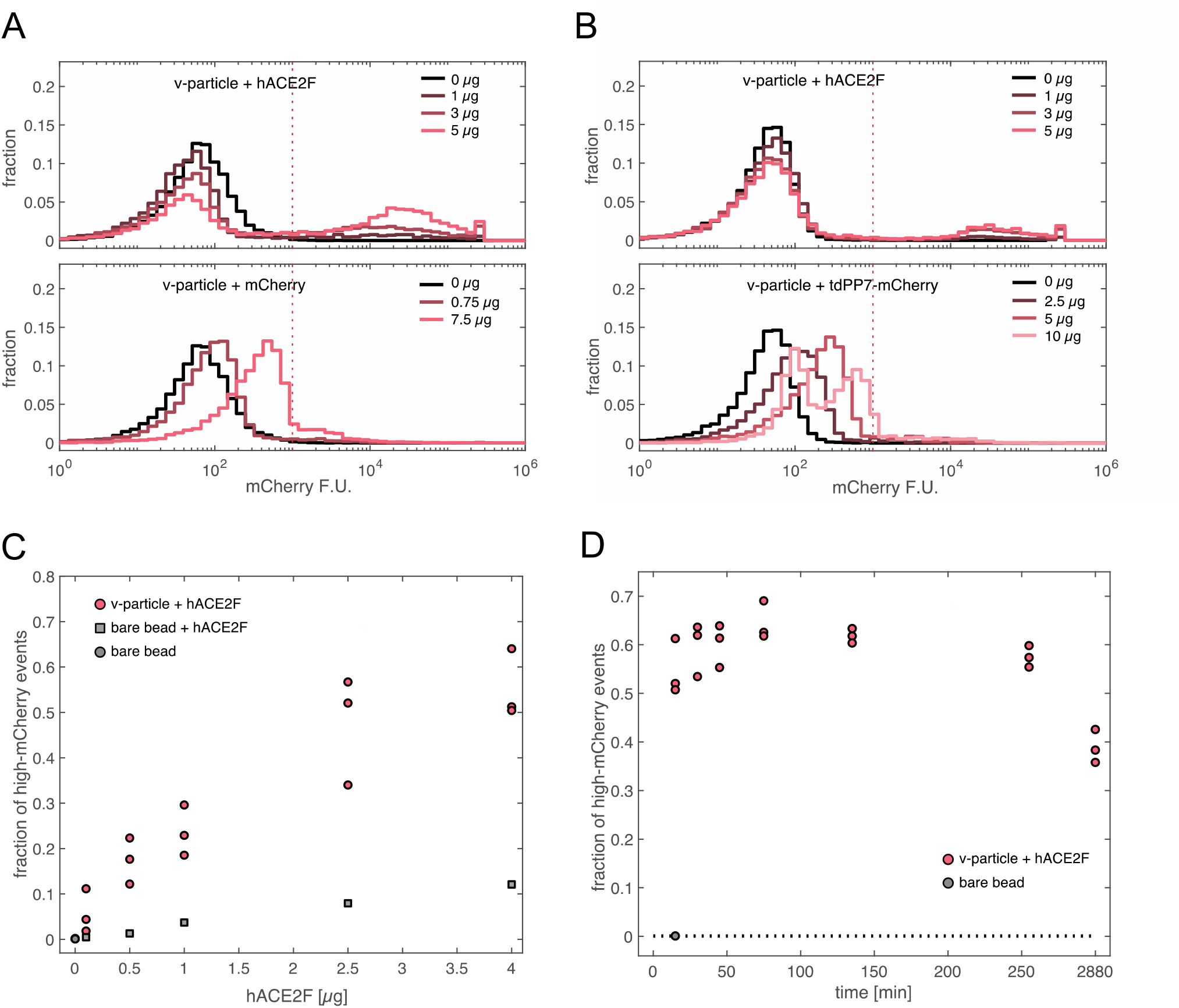
Optimizing v-particle assay using flow cytometry. (A) Flow cytometry assay results for v-particles with hACE2F and mCherry. (B) Flow cytometry assay results for 3-month-old v-particles with hACE2F and tdPP7-mCherry. (C) Sensitivity of v-particle - hACE2F binding. As low as ∼0.1 µg of hACE2F per reaction can be detected. (D) Fluorescence of v-particles as a function of reaction time. 45 min is sufficient for binding reactions.

To determine the sensitivity of v-particles to hACE2F, we measured the dependence of the v-particle fluorescence on hACE2F concentration. The results for the sensitivity assay are shown in Fig. 2C. We found that 2.5 µl of v-particles in our experimental conditions are sensitive to ∼0.1 µg of hACE2F, though a larger amount of hACE2F could provide more sensitivity when screening candidate inhibitors. We next determined the optimal time for binding reactions (Fig. 2D). Based on the results (Fig. 2D), we determined that 45 min is sufficient for binding reactions.

As a proof-of-concept for inhibitor screening, we measured the inhibition of v-particle - hACE2F binding in the presence and absence of a synthetic anti-RBD peptide, or sybody, Sb#68 ^28^. For details of Sb#68 expression see Supplementary Methods. We measured inhibition for different Sb#68 concentrations, in triplicate, as follows. The flow cytometry results for the different Sb#68 concentrations are plotted in Fig. 3A. For the v-particle reactions, we observe the high-fluorescence population indicating RBD-hACE2F binding (Fig. 3A-top). As the concentration of Sb#68 is increased, we see a dose-dependent reduction in the fraction of the highly fluorescent v-particles, indicating inhibition of the hACE2F interaction with the v-particles. We plot the fraction of the total particles in the high-fluorescence population as a function of the inhibitor (Sb#68) dose (Fig. 3B). The results show a continuous reduction is the high-population cell fraction, which provides a quantitative assay of RBD-hACE2F inhibition. For Sb#68, we get a 2-fold reduction in the fraction of bound hACE2F at 3.04 µg of inhibitor per 1 µl of v-particle stock.

**Figure 3.**
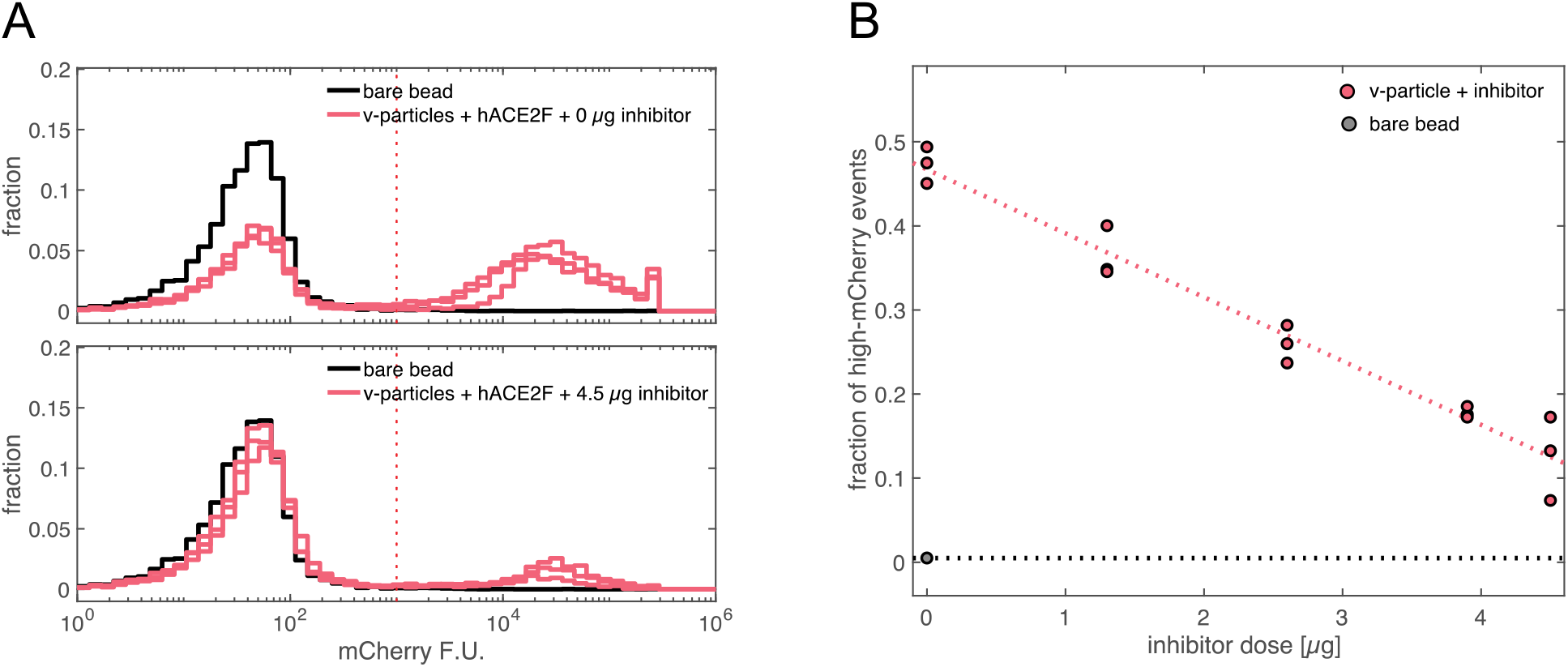
Inhibition of v-particle - hACE2F binding by Sb#68. (A) Flow cytometry data for 1 µl v-particle stock incubated with either 0 or 4.5 µg of Sb#68. (B) Percentage of v-particles with fluorescence above 1e3 as a function of inhibitor dose. Each Sb#68 concentration was measured in triplicate (using the same batch of v-particles). The fraction of v-particles in the high-mCherry fluorescence peak is reduced to half of the fraction in absence of inhibitor at a dose of 3.04 µg Sb#68 per 1 µl v-particle stock.

Finally, we utilized the v-particles to assess the efficacy of synthetic RNA-protein (SRNP) granules in binding to, and thereby depleting the active amount of, SARS-CoV-2 virions. Here, the v-particles provide a safe and microscopically-visible alternative to actual virions. RNA-protein granules can be produced in vitro ^29,30^, and have been shown to bind cellular components ^30,31^. We recently showed ^32^ that SRNP granules form specifically in vitro via self-assembly by mixing purified bacterial phage coat proteins with synthetic long non-coding RNA (slncRNA) molecules that encode multiple binding sites for the coat proteins. In this case, the protein component in the granule formulation was either tdPP7-mCherry, or hACE2F (see Supplementary Methods and Supplementary Table 1 for both proteins). The slncRNA component (slncRNA-PP7bsx14, see Supplementary Table 1 for sequence and Supplementary Methods for synthesis details) harbors 14 PP7 binding sites, to which the tdPP7 domain present in both hACE2F and tdPP7-mCherry can bind. The RNA thus increases the local concentration of hACE2F, which may facilitate virion entrapment and thus potentially function as an anti-SARS-CoV-2 decoy particle. To test for selective binding of the SRNP granules to the v-particles, we prepared the following samples: v-particles with slncRNA-PP7bsx14 and tdPP7-mCherry, v-particles with hACE2F, and v-particles with slncRNA-PP7bsx14 and hACE2F. We show the results of the binding experiments in Fig. 4. In the microscopy images, v-particles appear as green-fluorescent beads (Fig. 4 top-left). SRNP-granules appear as large red clumps or as bead-like particles (Fig. 4 bottom), which are located on the cover slip at different positions from the v-particles. When v-particles are mixed with hACE2F, colocalization of the hACE2F protein to the v-particles is observed, as expected from our previous experiments (Fig. 4 top-right). Finally, hACE2F-SRNP-granules appear to be bound to the v-particles (Fig. 4 bottom-right), as compared with the non-hACE2F-SRNP-granules which appear to be spatially separated from the v-particles (Fig. 4 bottom-left). Consequently, the SRNP-hACE2F granules provide a potential decoy or anti-SARS-CoV-2 therapeutic, which should be examined in follow-up research.

**Figure 4.**
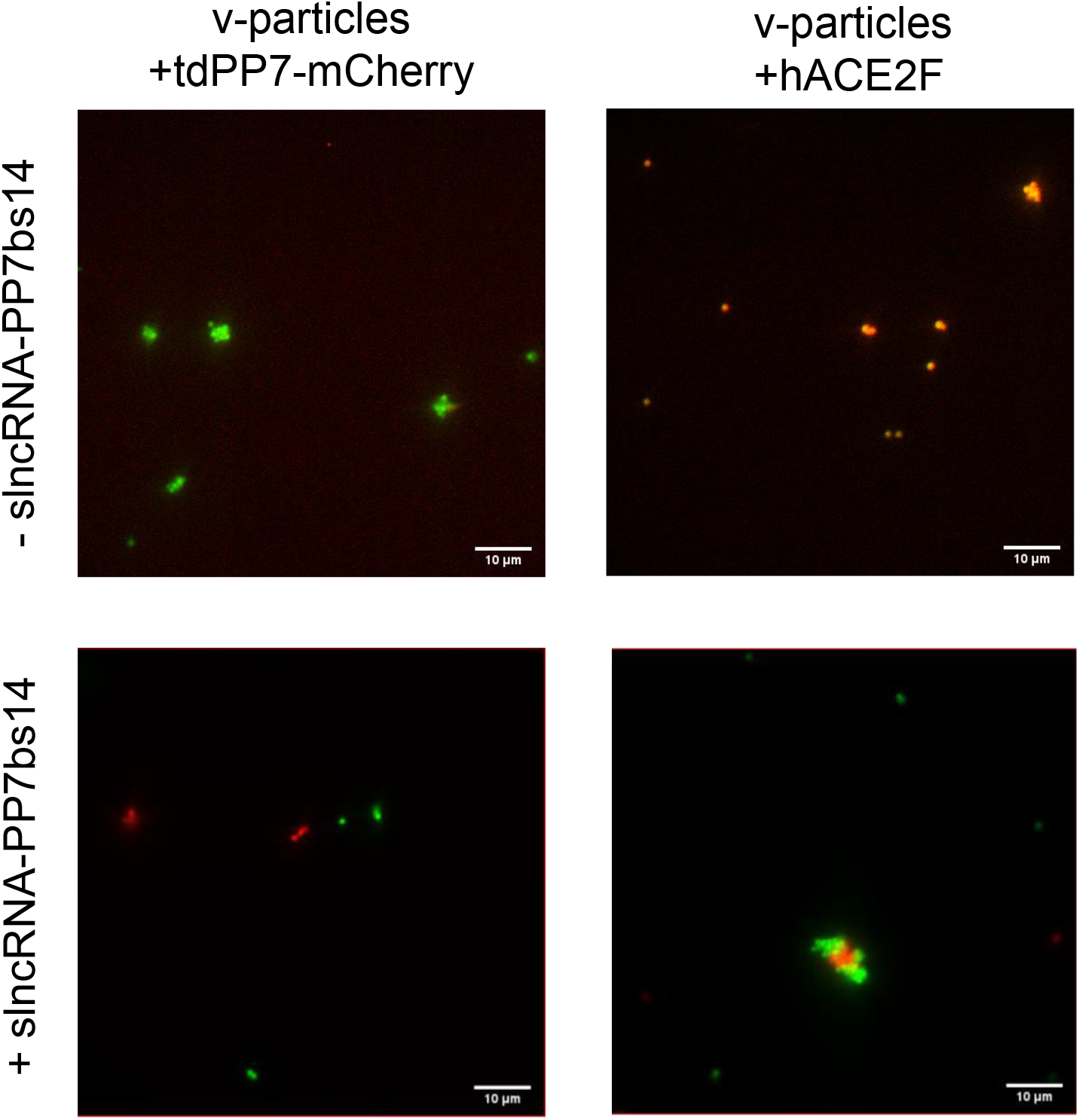
Entrapment of v-particles by slncRNA-PP7bsx14-hACE2F granules. Overlay of fluorescence microscopy images at 585 nm (mCherry) and 490 nm (FITC) excitation wavelengths, for (A) v-particles (from undiluted 1% w/v stock), (B) v-particles incubated with slncRNA-PP7bsx14 - tdPP7-mCherry granules, (C) v-particles incubated with hACE2F, and (D) v-particles incubated with slncRNA-PP7bsx14 - hACE2F granules. For (B-D), v-particle concentration was 0.1 % w/v. Protein concentrations in imaged samples were (B) 842 nM, (C) 560 nM, and (D) 507 nM. slncRNA-PP7bsx14 concentration in imaged samples was 112.8 nM (B, D).

We have presented a particle-based assay that enables rapid, cell-free screening of candidate inhibitors of protein-protein interaction, focusing on the interaction between SARS-CoV-2 Spike RBD bound to fluorescent particles (v-particles), and fluorescently-tagged hACE2 (hACE2F). The assay materials are commercially available or relatively easy to prepare, and do not include antibody components. The main difficulty in assay preparation is the production of the protein components. Depending on available lab resources, researchers may choose to outsource this step. We demonstrated the utility of the assay for quantifying inhibition of RBD-hACE2 interaction by the reported inhibitor Sb#68, as well as with a potential anti-SARS-CoV-2 RNP-granule decoy particle. Interestingly, the v-particles mixed with hACE2F exhibit a bi-modal or digital signal. Importantly for our assay, the signal from the high-expressing subpopulation is apparently highly sensitive to inhibitor concentration. The bi-modal signal suggests that hACE2F binding to v-particles is cooperative. This cooperativity may be due to steric displacement of BSA by tightly-bound hACE2F which promotes further binding of other hACE2F molecules, or by conformational change in bound hACE2F that results in stronger interaction with other hACE2F molecules. Although we described applications specific to RBD and hACE2F interaction, the presented applications could easily be modified to quantify interaction of other peptide-receptor interaction partners, such as RBD mutants with either hACE2 or other suspected host receptors ^33^, or other viral proteins with their respective host partners ^34^. We further demonstrated that v-particles can provide a cell-free alternative to more expensive and higher-biosafety-level cell-based assays for assessing proposed SARS-CoV-2 entrapment products. We hope that the relatively straightforward preparation, ease of use, and quantitative results of our v-particles and binding assay will have a significant impact in assays involving SARS-CoV-2 variants, as well as other viruses.

## MATERIALS AND METHODS

Details of protein expression and purification for his-tagged RBD, hACE2F, mCherry, and tdPP7-mCherry, appear in the Supplementary Information. Details of v-particle preparation also appear in the Supplementary Information.

### Flow-cytometry binding assays

v-particles and protein components (either binding partner hACE2F, or mCherry or tdPP7-mCherry as non-specific binding partner controls) were added according to the details below. Bovine serum albumin (BSA, 20 mg/ml, New England Biolabs) was added at a ratio of 5 -10 µg BSA per 1 µl of v-particle stock to all binding reactions to suppress non-specific binding of protein to the v-particles. Unless stated otherwise, samples were incubated at 37 °C with 145 rpm horizontal shaking for 45 min, covered in aluminum foil. All sample volumes were adjusted to 100 µl with 1x PBS + 10 µM ZnCl_2_ and measured via flow cytometry (MACSquant VYB, Miltenyi Biotec). The flow cytometer was calibrated using MacsQuant calibration beads (Miltenyi Biotec) before measurement, and 2 µl of 1% w/v amine polystyrene fluorescent yellow particles (NH_2_-beads, Spherotech, Inc) or carboxyl polystyrene fluorescent yellow particles (Spherotech, Inc.) in 100 µl 1x PBS were run as a negative control.

### Specificity of v-particle binding to hACE2F

In a Lo-Bind microcentrifuge tube, we combined 2.5 µl of pre-sonicated v-particle stock, 15 µg of BSA and either 1, 3, or 5 µg hACE2F, or 0.75 or 7.5 µg mCherry. Sample volumes were adjusted to 14 µl with 1x PBS + 10 µM ZnCl_2_. Samples were incubated, diluted, and measured by flow cytometry as described above.

### Sensitivity of v-particles to hACE2F

In a Lo-Bind microcentrifuge tube, 2.5 µl of pre-sonicated v-particle stock and 15 µg of BSA were mixed with one of the following amounts of hACE2F: 0, 0.1, 0.5, 1, 2.5, and 4 µg. The volume was adjusted to 9.5 µl with 1x PBS + 10 µM ZnCl_2_. For the negative control, v-particles were replaced with NH_2_-beads. Samples were incubated, diluted, and measured by flow cytometry as described above.

### Optimal time for binding reactions

In a Lo-Bind microcentrifuge tube, 1 µl of pre-sonicated v-particle stock, 10 µg of BSA, and 2 µg of hACE2F were added, and the volume was adjusted to 5.5 µl with 1x PBS + 10 µM ZnCl_2_. The samples were incubated at 37 °C with 145 rpm horizontal shaking for different amounts of time: 15, 30, 45, 75, 135, and 255 min, and 48 hr, covered in aluminum foil. Samples were diluted and measured by flow cytometry as described above.

### Inhibition of v-particle - hACE2F binding

In a Lo-Bind microcentrifuge tube, 1 µl of pre-sonicated v-particle stock, 10 µg of BSA, and Sb#68 in one of the following amounts: 0, 1.3, 2.6, 3.9, 4.5 µg were added, and the volume was adjusted to 5 µl with 1x PBS + 10 µM ZnCl_2_. Next, 5 µg of hACE2F was added to all the samples. This amount, chosen for convenience, was the highest amount we could use without further concentration of our stock. Samples were incubated, diluted, and measured by flow cytometry as described above.

### Selective binding of the SRNP granules to the v-particles

SRNP experiments were performed in granule buffer (GB: 750 mM NaCl, 1 mM MgCl_2_, 10% PEG 4000, in water). Reactions containing 8 µl GB, 1 µg tdPP7-mCherry or 1.5 µg hACE2F (in 1 µl), 0 or 1 µg slncRNA-PP7bsx14 (in 1 µl), and 0.5 µl Ribolock RNase Inhibitor (Thermo Fisher Scientific) were incubated at room temperature for 1 hr. After 1 hr, 1 µl from each reaction was deposited on a glass slide, together with 1 µl of pre-sonicated 1% w/v v-particle stock diluted 1:5 in water. A 1 µl control sample of undiluted v-particle stock was also deposited. After 10 minutes, the samples were sealed with coverslips and imaged using a 100x oil immersion objective on a Nikon Eclipse Ti epifluorescent microscope with iXon Ultra EMCCD camera (Andor) and NIS-Elements software (Nikon), with 585 nm (mCherry) and 490 nm (FITC) excitation using a CooLED PE illumination system (Andover).

## SUPPORTING INFORMATION

The following files are available free of charge.

A list of the nucleotide sequences of all proteins used in the described assay (Supplementary Table S1, CSV). A description of methods used for expression and extraction of the proteins used in this work, and a description of v-particle generation (Supplementary Methods, PDF).

## AUTHOR INFORMATION

### Author Contributions

N. Kikuchi, O. W., N. G., S. G., and R. A. conceived the approach and designed the experiments. N. Kikuchi prepared the v-particles and carried out binding experiments. N. G. prepared slncRNA-PP7bsx14 and carried out microscopy experiments. O. W. and S. G. expressed and purified RBD, hACE2F, mCherry, and tdPP7-mCherry, and assisted with binding experiments. N. N. assisted with RBD secretion and purification. N. Katz assisted with slncRNA-PP7bsx14 design. S. A., E. S., and T. A. designed the Sb#68 expression vector. S. A. expressed and purified Sb#68. N. Kikuchi, O. W., N. G., S. G., and R. A. prepared the manuscript. All authors have given approval to the final version of the manuscript.

## NOTES

N. Kikuchi, O. W., N. G., S. G., and R. A. are inventors on US Provisional Patent Application No. 63/187969 concerning some of the described technologies. N. Katz and R. A. are inventors on US Patent Application 2021/0095296 A1.

## ACKNOWLEDGEMENT

We thank Omer Yehezkeli, Smadar Shulami, Onit Alalouf, and Moran Bercovici from the Technion for expert advice. N. Kikuchi acknowledges the support of the Zuckerman STEM Leadership program. We thank J. D. Walter for advice regarding sybodies. We thank Integrated DNA Technologies for contributing the gBlock encoding Sb#68, and Promega for contributing the *E. coli* KRX cells, to the Technion 2020 iGEM team.

## ABBREVIATIONS

RBD: SARS-CoV-2 Spike receptor-binding domain
hACE2: human angiotensin-converting enzyme 2
tdPP7: tandem-dimer form of bacteriophage PP7 coat protein
hACE2F: fluorescently-labeled hACE2 also containing tdPP7
slncRNA-PP7bsx14: synthetic long noncoding RNA harboring 14 binding sites of bacteriophage PP7 coat-protein

## REFERENCES

1. Zhu, N. et al. A Novel Coronavirus from Patients with Pneumonia in China, 2019. N. Engl. J. Med. (2020) doi:10.1056/NEJMoa2001017.

2. Zhou, P. et al. A pneumonia outbreak associated with a new coronavirus of probable bat origin. Nature 579, 270–273 (2020).

3. Yan, R. et al. Structural basis for the recognition of SARS-CoV-2 by full-length human ACE2. Science 367, 1444–1448 (2020).

4. Wang, Q. et al. Structural and Functional Basis of SARS-CoV-2 Entry by Using Human ACE2. Cell 181, 894-904.e9 (2020).

5. Rees-Spear, C. et al. The impact of Spike mutations on SARS-CoV-2 neutralization. bioRxiv 2021.01.15.426849 (2021) doi:10.1101/2021.01.15.426849.

6. Krammer, F. SARS-CoV-2 vaccines in development. Nature 586, 516–527 (2020).

7. Forni, G. & Mantovani, A. COVID-19 vaccines: where we stand and challenges ahead. Cell Death Differ. 28, 626–639 (2021).

8. Zhang, Y. et al. Safety, tolerability, and immunogenicity of an inactivated SARS-CoV-2 vaccine in healthy adults aged 18–59 years: a randomised, double-blind, placebo-controlled, phase 1/2 clinical trial. Lancet Infect. Dis. 21, 181–192 (2021).

9. Voysey, M. et al. Safety and efficacy of the ChAdOx1 nCoV-19 vaccine (AZD1222) against SARS-CoV-2: an interim analysis of four randomised controlled trials in Brazil, South Africa, and the UK. The Lancet 397, 99–111 (2021).

10. Polack, F. P. et al. Safety and Efficacy of the BNT162b2 mRNA Covid-19 Vaccine. N. Engl. J. Med. 383, 2603–2615 (2020).

11. Baden, L. R. et al. Efficacy and Safety of the mRNA-1273 SARS-CoV-2 Vaccine. N. Engl. J. Med. 384, 403–416 (2021).

12. Formica, N. et al. Evaluation of a SARS-CoV-2 Vaccine NVX-CoV2373 in Younger and Older Adults. medRxiv 2021.02.26.21252482 (2021) doi:10.1101/2021.02.26.21252482.

13. Huang, Y., Yang, C., Xu, X., Xu, W. & Liu, S. Structural and functional properties of SARS-CoV-2 spike protein: potential antivirus drug development for COVID-19. Acta Pharmacol. Sin. 41, 1141–1149 (2020).

14. Hoffmann, M. et al. SARS-CoV-2 Cell Entry Depends on ACE2 and TMPRSS2 and Is Blocked by a Clinically Proven Protease Inhibitor. Cell 181, 271-280.e8 (2020).

15. Pushpakom, S. et al. Drug repurposing: progress, challenges and recommendations. Nat. Rev. Drug Discov. 18, 41–58 (2019).

16. Mercorelli, B., Palù, G. & Loregian, A. Drug Repurposing for Viral Infectious Diseases: How Far Are We? Trends Microbiol. 26, 865–876 (2018).

17. Galindez, G. et al. Lessons from the COVID-19 pandemic for advancing computational drug repurposing strategies. Nat. Comput. Sci. 1, 33–41 (2021).

18. Kuleshov, M. V. et al. The COVID-19 Drug and Gene Set Library. Patterns 1, 100090 (2020).

19. Nie, J. et al. Quantification of SARS-CoV-2 neutralizing antibody by a pseudotyped virus-based assay. Nat. Protoc. 15, 3699–3715 (2020).

20. Neerukonda, S. N. et al. Establishment of a well-characterized SARS-CoV-2 lentiviral pseudovirus neutralization assay using 293T cells with stable expression of ACE2 and TMPRSS2. PLOS ONE 16, e0248348 (2021).

21. Contreras-Llano, L. E. & Tan, C. High-throughput screening of biomolecules using cell-free gene expression systems. Synth. Biol. 3, (2018).

22. Medhi, R., Srinoi, P., Ngo, N., Tran, H.-V. & Lee, T. R. Nanoparticle-Based Strategies to Combat COVID-19. ACS Appl. Nano Mater. 3, 8557–8580 (2020).

23. Norman, M. et al. Ultrasensitive high-resolution profiling of early seroconversion in patients with COVID-19. Nat. Biomed. Eng. 4, 1180–1187 (2020).

24. Dogan, M. et al. SARS-CoV-2 specific antibody and neutralization assays reveal the wide range of the humoral immune response to virus. Commun. Biol. 4, 1–13 (2021).

25. Baker, M. How quality control could save your science. Nature 529, 456–458 (2016).

26. Amanat, F. et al. A serological assay to detect SARS-CoV-2 seroconversion in humans. Nat. Med. 1–4 (2020) doi:10.1038/s41591-020-0913-5.

27. Watanabe, Y., Allen, J. D., Wrapp, D., McLellan, J. S. & Crispin, M. Site-specific glycan analysis of the SARS-CoV-2 spike. Science (2020) doi:10.1126/science.abb9983.

28. Walter, J. D. et al. Sybodies targeting the SARS-CoV-2 receptor-binding domain. bioRxiv 2020.04.16.045419 (2020) doi:10.1101/2020.04.16.045419.

29. Lin, Y., Protter, D. S. W., Rosen, M. K. & Parker, R. Formation and Maturation of Phase-Separated Liquid Droplets by RNA-Binding Proteins. Mol. Cell 60, 208–219 (2015).

30. Nakamura, H. et al. Intracellular production of hydrogels and synthetic RNA granules by multivalent molecular interactions. Nat. Mater. 17, 79–89 (2018).

31. Han, T. W. et al. Cell-free Formation of RNA Granules: Bound RNAs Identify Features and Components of Cellular Assemblies. Cell 149, 768–779 (2012).

32. Granik, N., Katz, N., Goldberg, S. & Amit, R. Formation of synthetic RNP granules using engineered phage-coat-protein - RNA complexes. bioRxiv 682518 (2021) doi:10.1101/682518.

33. Daly, J. L. et al. Neuropilin-1 is a host factor for SARS-CoV-2 infection. Science 370, 861– 865 (2020).

34. Li, F. Structure, Function, and Evolution of Coronavirus Spike Proteins. Annu. Rev. Virol. 3, 237–261 (2016).

