## Supplementary methods for "A cell-free assay for rapid screening of inhibitors of hACE2-receptor - SARS-CoV-2-Spike binding"

**Supplementary Methods for paper entitled ``A cell-free assay for rapid screening of inhibitors of hACE2-receptor - SARS-CoV-2-Spike binding''**

**RBD mammalian expression and purification:** The plasmid encoding RBD was a gift from the Krammer lab. The plasmid was transformed into *E. coli* TOP10 cells (Invitrogen) and miniprepped (ZymoPure plasmid miniprep II, Zymo). 293F cells were cultured in 30 ml Freestyle 293 [supplemented with penicillin-streptomycin solution (Biological Industries) at 0.5% v/v] expression medium (Thermo Fisher), in 125 ml flat-bottom flasks (TriForest), at 37 °C with 8% $CO_2$ and 135 rpm shaking. 24h before transfection, cells were passed at 0.6-0.7e6 cells/mL and grown overnight. On the day of transfections, cells were diluted to 1e6/ml cell concentration and were then transfected as follows: 37.5 µg plasmid DNA and 120 µl of 0.5 mg/ml branched polyethylenimine (PEI, MW ~25,000, Sigma Aldrich) were separately brought to 600 µl in Opti-MEM (Gibco), and incubated for 5 min. PEI solution was added to DNA solution and incubated at room temperature for 15 min. 1200 µl PEI+DNA solution was added to the 30 ml culture. After 5-6 days of incubation at 37 °C with 8% $CO_2$ at 135 rpm shaking, cells were centrifuged for 5 min at 100xg, supernatant containing secreted his-tagged RBD was collected, and cells were discarded. The RBD-containing supernatant was incubated with Ni-coated beads (either Purecube 100 Indigo, Cube Biotech, or Hislink protein purification resin, Promega) at room temperature for 1 hr, with 13 rpm overhead rotation. The his-tagged proteins were then purified on a gravity-flow column (PolyPrep chromotography column, Biorad). In our hands, the elution buffer from the Cube protocol (EB: 50mM $NaH_2PO_4$, 300 mM NaCl, and 500 mM imidazole in deionized water, pH 8.0) worked better for both types of Ni-coated beads. Typical RBD yield was ~1 mg from 90-120 mL of 293F culture. We changed the buffer of the eluted RBD to phosphate buffered saline (PBS: Dulbecco's phosphate buffered saline -calcium -magnesium, Biological Industries) by rinsing multiple times with 1x PBS on a 3 kDa MWCO spin column (Amicon Ultra 0.5 mL, Merck Millipore). RBD was stored at -20 °C. Lengths of RBD and all other proteins in this work were verified by SDS polyacrylamide gel (SDS-PAGE) followed by Coomassie staining.

**hACE2-mCherry-tdPP7 (hACE2F) mammalian expression and purification:** The plasmid encoding the extracellular domain of ACE2 (amino acids 18 to 740) fused to tdPP7 (hACE2-tdPP7) with C-terminal his tag was ordered from Twist Bioscience (using different coding sequences for the two copies of PP7 coat protein), and modified in the lab to add mCherry (see full sequence in Supplementary Table 1). The transfection, growth, expression, and purification were similar to RBD expression. Typical hACE2F yield was ~1 mg from 90-120 mL of 293F culture. The culture, supernatant, and Ni-coated beads were visibly pale pink during expression and purification stages. After elution, the 2-3 mL hACE2F sample was dialyzed twice against 800 mL of 1x PBS + 10 µM $ZnCl_2$ (Pur-A-Lyzer Maxi 3500 dialysis unit, Sigma Aldrich), further concentrated on a 3 kDa MWCO spin column (Amicon Ultra 0.5 mL, Merck Millipore), and stored at -20 °C.

**mCherry bacterial expression and purification:** A bacterial plasmid encoding his-tagged mCherry under the rhlR promoter (containing the las box, inducible by N-butyryl-L-Homoserine lactone [C4-HSL], Cayman Chemical), ampicillin resistance, and RhlR was transformed into *E. coli* TOP10 cells (Invitrogen). Cells containing the plasmid were grown in 10 ml Luria-Bertani medium (LB: 10 g NaCl, 10 g tryptone, and 5 g yeast extract in 1 L deionized water, autoclaved) containing 100 µg/ml ampicillin (Amp) in a 50 ml falcon overnight, at 37 °C and 250 rpm. The culture was diluted into 500 ml terrific broth (TB: 24 g yeast extract, 20 g tryptone, 4 ml glycerol in 1 L of water, autoclaved, and supplemented with 17 mM $KH_2PO_4$ and 72 mM $K_2HPO_4$) containing 100 µg/ml Amp and 97 nM C4-HSL in a 2-liter flask, and grown for another day at 37 °C and 250 rpm. Culture was visibly pink the next morning. Cells were centrifuged at 8000 rpm for 10 min in 250 ml bottles, supernatant was discarded, and the visibly pink pellets were resuspended in resuspension buffer (RB: 50 mM Tris, 100 mM NaCl, 0.02% sodium azide in deionized water, pH 7.0). The resuspended cells were lysed by passing the culture four times through a high-pressure homogenizer (Emulsiflex, Avestin Inc, Canada) at an average working pressure of 10-15 kpsi and maintained at 4 °C using a circulating bath (GMBH, Germany). Collected lysate was centrifuged at 13 krpm for 30 min. Clear, visibly pink supernatant was collected, and cell debris was discarded. Typical mCherry yield was 10 mg from 500 mL of TB culture. mCherry buffer was changed by rinsing multiple times with 1x PBS on a 3 kDa MWCO spin column, and mCherry was stored at -20 °C.

**tdPP7-mCherry bacterial expression and purification:** See details for mCherry expression and extraction, with mCherry replaced by mCherry-tdPP7 (see Supplementary Table 1 for sequence).

**Sb#68 bacterial expression and purification:** We expressed his-tagged Sb#68 (see Supplementary Table S1 for sequence, ordered as a gBlock from Integrated DNA Technologies, IDT) from a pET9D bacterial plasmid under a T7 promoter, in *E. coli* KRX cells (Promega). Growth and expression were similar to mCherry, only with 25 µg/ml kanamycin instead of Amp, and with 0.1% w/v rhamnose instead of C4-HSL for induction. Extraction and buffer change to 1xPBS were the same as described earlier for mCherry. Sb#68 yield was ~5 mg from 500 mL of TB culture.

**Generation of v-particles:** SPHERO carboxyl fluorescent yellow particles with 0.7-0.9 µm diameter (Spherotech Inc., specified batch diameter was 0.92 µm) were sonicated in their original container for 3 min, with multiple vortex mixing. 100 µl of 1% w/v particles were transferred into a Lo-Bind microcentrifudge tube (Eppendorf) and centrifuged for 15 min at 3000xg. The supernatant was removed and 100 µl of 50 mM MES buffer was added [MES stock: 0.5 M 2-(N-Morpholino) ethanesulfonic acid (Sigma Aldrich) in deionized water, at pH5; diluted to 50 mM in deionized water]. The sample was vortexed until particle aggregation was not visible and the mixture looked "milky". The sample was centrifuged again for 15 min at 3000xg and the supernatant was replaced with 50 µl of 50 mM MES containing 0.1 mg N-(3-Dimethylaminopropyl)-N′-ethylcarbodiimide hydrochloride (EDC, Sigma Aldrich) and 50 µl of 50 mM MES containing 1.1 mg N-hydroxysulfosuccinimide sodium salt (Sulfo-NHS, Sigma Aldrich). The sample was vortexed and incubated at room temperature with 145 rpm horizontal shaking for 30 min covered in aluminum foil. The sample was then centrifuged for 15 min at 3000xg and the supernatant was replaced with 100 µl of 1x PBS, 2 times. The sample centrifuged again for 15 min at 3000xg and the supernatant was replaced with 30 µg of RBD in 100 µl of 1x PBS, and incubated at room temperature with 145 rpm horizontal shaking for 2.5 hrs covered in aluminum foil. The sample was centrifuged for 15 min at 3000xg and the supernatant was replaced with 100 µl of 1x PBS + 10 µM $ZnCl_2$, 3 times. The synthesized v-particle stock was stored at 4 °C. Final fluorescent particle concentration in the v-particle stock is approximately 1% w/v. The number of particles in 1 mL is approximately 35e9 for 0.8 µm particles at 1% w/v (https://www.spherotech.com/particle.html). The maximum covalent attachment ratio of RBD to the particles is 50 µeq/g (equal to the manufacturer's claim of 50 µeq/g carboxyl groups). This yields a maximum ratio of approximately 3e5 RBD per particle, based solely on the number of available functional groups. The actual ratio is likely lower due to partial binding, protein size, and steric effects.

**slncRNA-PP7bsx14 synthesis:** DNA encoding a T7 promoter (TAATACGACTCACTATA with trailing GGG) followed by 14 non-repetitive binding sites of bacteriophage PP7 coat protein with EcoRI (and unused NruI) restriction sites on both ends was ordered as a gBlock (IDT) (see Supplementary Table 1 for sequence), cloned into a pCMV cloning vector in *E. coli* TOP10 (Lucigen) using the EcoRI sites, miniprepped (NucleoSpin Plasmid Mini, Macherey-Nagel), restricted with EcoRI (New England Biolabs, NEB), and column-cleaned (Wizard SV Gel and PCR Clean-Up System, Promega). slncRNA-PP7bsx14 was transcribed in vitro from the resulting DNA in a 30 µl reaction at 37 °C for 3 hours (HiScribe T7 High Yield RNA Synthesis Kit, NEB). The reaction volume of the transcription product was diluted to 90 µl using UltraPure water (Bio-Lab Ltd.), 10 µl of DNAse I buffer and 2 µl of DNAse I (NEB) were added, and the resulting mix was incubated at 37 °C for 15 min. Finally, slncRNA-PP7bsx14 was purified (Monarch RNA Cleanup Kit 500 µg, NEB), and stored for later use at -80 °C. Typical concentrations were 100-1000 ng/µl, with 100 µl final volume.
